# Regionally specific resting-state beta neural power predicts brain injury and symptom recovery in adolescents with concussion: a longitudinal study

**DOI:** 10.1101/2025.01.22.633765

**Authors:** J. Christopher Edgar, Lisa Blaskey, Yuhan Chen, Olivia E. Podolak, Drayton L. Murray, Marybeth McNamee, Kimberly Konka, Jeffrey I. Berman, Timothy P.L. Roberts, Mingxiong Huang, Kristy B. Arbogast, Christina L. Master

## Abstract

Mild traumatic brain injury (mTBI) is common in adolescents. Magnetoencephalography (MEG) studies (primarily reporting on adult males) have demonstrated abnormal resting-state (RS) brain activity in mTBI. The present study sought to identify RS abnormalities in male and female adolescents with mTBI (no previous mTBI and no previous DSM-5 diagnosis) identified from an outpatient specialty care concussion program setting as a basis for evaluating potential clinical utility. Visit 1 MEG RS data were obtained from 46 adolescents with mTBI (mean age: 15.4 years, 25 females) within 4 months of a mTBI (mTBI acute to sub-acute period) as well as from 34 typically developing (TD) controls (mean age: 14.8 years; 17 females) identified from the local community. Visit 2 RS data (follow-up ∼4.3 months after Visit 1; mTBI sub-chronic period) were obtained from 36 mTBI (19 females) and 29 TD (14 females) of those participants. Source-space RS neural activity was examined from 4 to 56 Hz. Visit 1 t-tests showed that group differences were largest in the beta range (16-30 Hz; mTBI < TD), with Visit 2 whole-brain linear mixed model (LMM) analyses examining beta-band group differences as a function of Visit. A main effect of Group indicated Visit 1 and 2 beta-band group differences in midline superior frontal gyrus, right temporal pole, and right central sulcus (all mTBI < TD). The group effects were large (Cohen’s *d* values 0.75 to 1.31). Of clinical significance in the mTBI group, a decrease in mTBI symptoms from Visit 1 to 2 was associated with an increase in beta power in 4 other brain regions. Present findings suggest that RS beta power has potential as a measure and perhaps as a mechanism of clinical recovery in adolescents with mTBI.

## Background

Mild traumatic brain injury (mTBI; also referred to as concussion) results from biomechanical forces that impart acceleration and deceleration to the head and result in a temporary loss of normal brain function. The United States Centers for Disease Control and Prevention estimates that up to 3.6 million mTBI occur annually in the United States (1, 2). The majority of these injuries occur in youth 5-18 years of age, with those 11-18 years of age representing the largest proportion of those injured (2). The cause of youth mTBI is varied and includes motor vehicle crashes, sport-related injuries, falls, and assaults. These mechanisms of injury result in clinical manifestations such as physical signs and symptoms (eyes movement deficits, headache, dizziness, nausea), cognitive deficits (memory, attention, executive functioning), and emotional issues (irritability, sadness, anxiety, depression) (3–7). Adolescents may be more vulnerable to the long-term effects of brain injury, as they recover more slowly than adults (8–10).

mTBI is primarily a dichotomous clinical diagnosis, relying on a patient’s subjective report of symptoms (11) as well as qualitative and quantitative clinical assessments that have effort-dependent components and/or are influenced by learning effects (12). A limitation of this diagnostic tradition is that it provides no direct information regarding which brain regions are affected or the anatomical or physiological features of the brain injury. To address this gap, objective diagnostics are needed to identify the brain changes associated with mTBI and to provide a method to track changes in brain pathology and recovery across time (13, 14).

Clinical brain imaging based on lesion detection or structural abnormality has not been found to be useful for identifying mTBI (15–19). For example, for mTBI, intracranial lesions were observed with conventional neuroimaging (MRI and CT) in only 4%, 16%, and 28% of patients with Glasgow Coma Scale scores of 15, 14, and 13, respectively (20). The structural damage in mTBI is described as traumatic axonal injury – with microscopic white-matter injury thought to be an important hallmark of structural brain pathology (14). Standard diffusion tensor imaging (DTI) clinical interpretation, however, does not detect mTBI in the acute phase or at the individual patient level (14, 21).

Several adult studies indicate that the assessment of neural brain activity via magnetoencephalography (MEG) shows promise in identifying abnormal brain activity in individuals with mTBI. MEG is a non-invasive functional imaging technique that measures the neuronal current in gray matter with high temporal resolution (<1 ms) and good spatial localization accuracy (2–3 mm at cortical level) (22). Seminal studies from Lewine et al. (23, 24) showed that MEG is sensitive to abnormal resting-state (RS) slow-wave activity (1–6 Hz) in adults with persistent (> 1 year) post-concussive symptoms following mTBI.

Since the reports by Lewine et al. (23, 24), other MEG studies have reported atypical RS neural activity in adults with mTBI. Allen et al. (25) reviewed nearly 40 MEG mTBI studies conducted since 2021, reporting on the pattern of RS findings as well as findings observed from working memory and attention tasks (for a similar review, see Peitz et al. (26)). Allen et al. noted that, across MEG RS studies, the most common finding was higher power in the delta frequency band in adults with mTBI versus controls (8 of the 14 studies), with group differences most often localized to the frontal, temporal, and parietal lobes. Group differences in other frequency bands were also noted, including more RS alpha power (27–30), less beta power (31), and more gamma power in individuals with mTBI than controls (32), with several studies observing widespread connectivity changes across several frequency bands (29, 33–35). They concluded their review by noting that although “MEG has demonstrated clear promise as a functional neuroimaging modality, it is not yet a diagnostic or prognostic clinical tool in mTBI of sufficient sensitivity and specificity. However, MEG is one of the most sensitive imaging modalities for the evaluation of mTBI, considering the very low sensitivity of CT, structural MRI and EEG” (p. 9). A similar conclusion was reached by a panel of mTBI experts convened to address the utility of MEG for identifying brain injury in individuals with mTBI by the United Kingdom Ministry of Defense (36).

Examination of the MEG mTBI studies reported in Allen et al. (25) shows that the majority included mostly males (46% included <u>only</u> male participants) and were cross-sectional. Recently (2020+), four MEG studies have reported on RS brain activity in pediatric mTBI populations (37–40). As detailed in the present Discussion, these pediatric studies (mostly male participants) observed mTBI / control group differences (power and functional connectivity) similar to those in adults. The present study sought to address gaps in the newly developing pediatric mTBI literature via a longitudinal study assessing RS brain activity in male and female adolescents with mTBI to determine whether findings in younger and older samples generalize to adolescents as a basis for evaluating potential clinical utility. Given RS electrophysiology abnormalities in individuals with depression, anxiety, and learning disorders such as attention-deficit/hyperactivity disorder (for a recent review see (41)), adolescents with mTBI with an anxiety, depression, or learning disorder diagnosis prior to the concussion were excluded so that the results could be more easily interpreted (see details in Section 2.1).

## Methods

### Participants and Clinical Assessments

Adolescents 12 to 18 years old with a mTBI were recruited from the Children’s Hospital of Philadelphia (CHOP) specialty care concussion program. Typically developing (TD) adolescents 12 to 18 years old were recruited from CHOP sports medicine specialty care practices, parent emails, and flyers posted around the Philadelphia metropolitan area. As shown in Figure 1, the study was designed for adolescents with mTBI to complete Visit 1 within 3 months following their concussion and Visit 2 ∼4 months later. As detailed below, due to the COVID-19 pandemic some of the Visit 1 and Visit 2 adolescents completed their exam outside the planned Visit 1 or 2 period.

**Figure 1.**
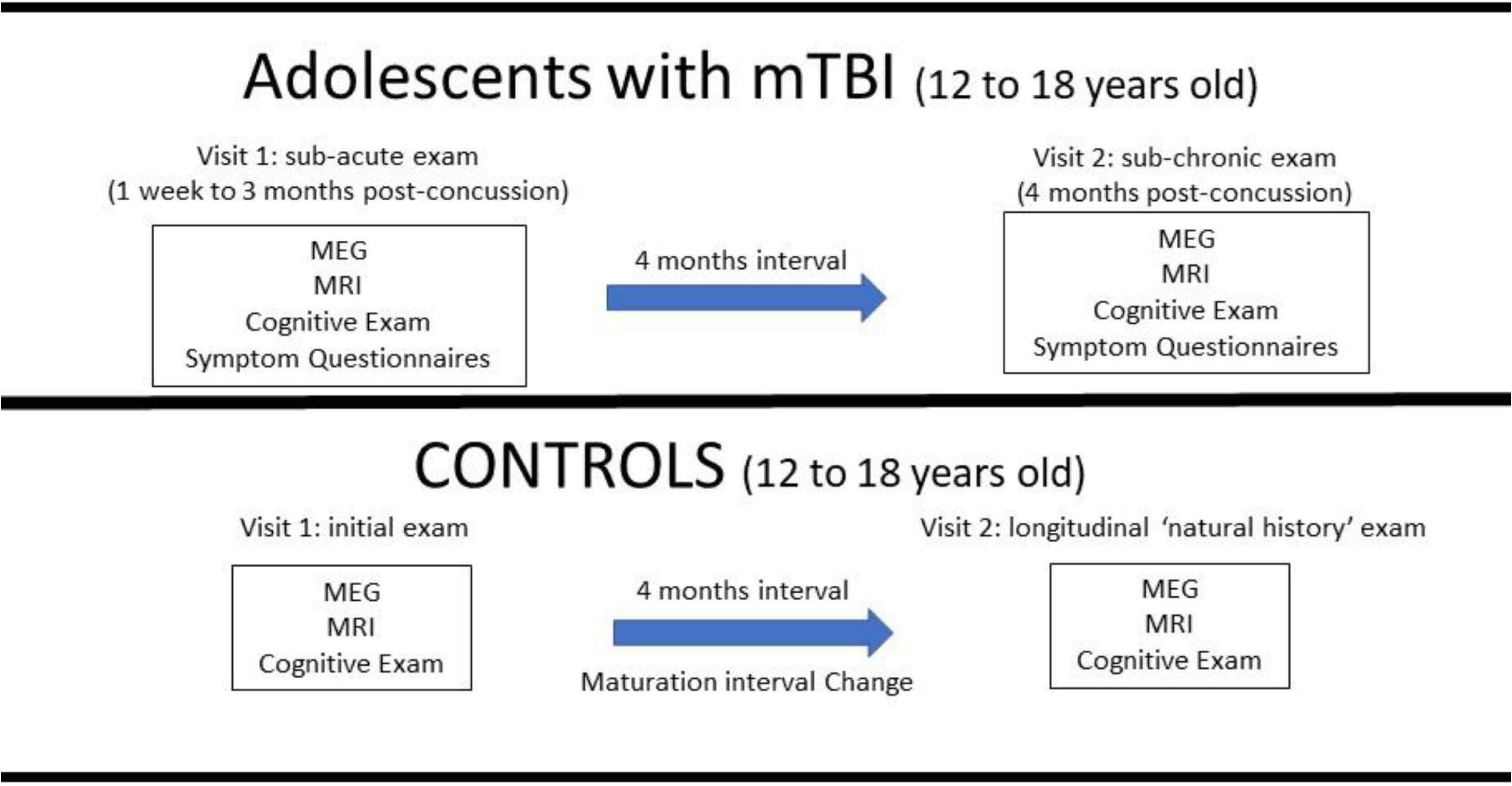
In the adolescents with mTBI, Visit 1 occurred within 4 months of their injury and Visit 2 ∼4 months later. In the controls, Visit 1 and 2 were separated by ∼4 months.

Adolescents with TBI and TD adolescents were selected according to the following criteria: a) English as primary/dominant language, b) no history of neurological, psychiatric, developmental, or neurogenetic disorders (based on parent screening interview, participant questionnaires (see below), and when possible a review of medical record), c) not pregnant, d) no premature birth (<34 weeks), e) no history of psychotropic medications, and f) no uncorrected vision or hearing impairments. Injured adolescents met the criteria for mTBI specified in the Consensus Statement on Concussion in Sport (7, 42) as diagnosed by our team. Exclusion criteria for adolescents with mTBI included head injury within one year prior to the presenting injury. Exclusion criteria for TD adolescents included any history of TBI.

During the phone screen at Visit 1 and 2, participants from both groups were evaluated to exclude those with DSM-5 diagnoses based on a review of the child’s medical history and results of parent ratings on a standardized behavior questionnaire (Child and Adolescent Symptom Inventory-5 (CASI-5, (43)). History of prior mental health diagnosis (e.g., mood or anxiety) and ongoing or prior (last 12 months) treatment was exclusionary at time of telephone screen, as were parent ratings in the clinically significant range on the CASI-5 (i.e., DSM-5 disorder-specific symptom count criteria exceeded diagnostic cut-offs and/or subdomain dimensional symptom severity standardized T-scores were 70 or higher (≥ 98^th^ percentile)). At the in-person Visit 1 and 2, additional standardized ratings provided further information to rule-out DSM-5 diagnoses as well as to assess symptoms in cognitive, emotional, sleep, and physical domains (7, 44), including the Youth (Self Report) Inventory - 4R (YI-4R) (43), the Child Behavior Checklist (CBCL parent report (45)), the Child Depression Inventory-2 (CDI-2, child self-report (46)), the Screen for Child Anxiety and Related Disorders (SCARED; child self-report (47, 48)), and the Behavior Rating Inventory of Executive Functioning (BRIEF; parent report (49))). The YI-4R is a DSM-5 reference rating scale that screens for emotional and behavioral symptoms among adolescents 12-18 years old (43), completed by the participant during the visit and used to identify participants with a likely current DSM psychiatric diagnosis. The CBCL provides an index of DSM-IV symptoms in adolescents 6-18 years old. The CDI-2 was administered to screen for childhood depression and the SCARED for childhood anxiety. The BRIEF is an 86-item survey administered to parents to inquire about their child’s executive functioning. Scores in the clinically significant range (standardized T-scores ≥ 70 = 98^th^ percentile or higher) were reviewed by the study pediatric psychologist (LB), and participants were excluded if there was clinical concern about the presence of a DSM-5 diagnosis.

Cognitive ability was assessed at Visit 1 and 2, with a focus on memory, processing speed, and other aspects of executive functioning (EF). The following tests were administered: Digit Span (attention/working memory), Coding, and Symbol Search (processing speed) from the Wechsler Intelligence Scale for Children-V/Wechsler Adult Intelligence Scale-IV ((50)), the Trail Making Test (EF-switching), Color-Word Interference Test (EF-interference control/inhibition), and Verbal Fluency Test (EF-fluency) from the Delis-Kaplan Executive Function System (DKEFS; (51)), and the Rey Auditory Verbal Learning Test (RAVLT) for memory (52). Alternate test versions were used at Visit 2 when such forms existed (e.g., for DKEFS Verbal Fluency and RAVLT).

Finally, to assess symptoms associated with a mTBI, at Visit 1 and 2 the adolescents with mTBI were administered the Post-Concussion Symptom Inventory (PCSI (44)), a questionnaire that assesses concussion symptoms in cognitive, emotional, sleep, and physical domains.

Due to the COVID-19 pandemic, some of the above clinical measures were not obtained from some participants, especially the cognitive measures. In particular, whereas most participants and one of their parents completed the symptoms questionnaires (e.g., the PCSI), cognitive measures were not obtained from many participants when our institution discouraged face-to-face research assessments early in the pandemic, and many families opted out. Given that only 10 participants with mTBI had both Visit 1 and 2 cognitive measures, analyses focused only on PCSI Total score.

### MEG and Structural MRI Data Acquisition

Eyes-closed RS MEG data were obtained from adolescents with mTBI and TD using a 275-channel MEG system (VSM MedTech Inc., Coquitlam, BC). Adolescents were instructed to close their eyes for 5 minutes, with three separate 5-minute recordings obtained. Electro-oculogram (EOG) (vertical EOG above and below the left eye) and electrocardiogram (ECG) were also obtained. During the recording, the EOG channel was monitored, and, if the participant opened their eyes during the recording, they were reminded to close them. After applying a band-pass filter (0.03 to 150 Hz), EOG, ECG, and MEG signals were digitized at 1,200 Hz with 3rd-order gradiometer environmental noise reduction. Structural MRI (sMRI) provided T1-weighted, 3-D MPRAGE anatomical images for source localization (3T Siemens Prisma scanner). To assess for brain injury pathology visible on conventional MR, T1-weighted images, susceptibility weighted images (SWI), T2 FLAIR, and T2-weighted images were obtained. sMRI scans were clinically reviewed by a pediatric neuroradiologist.

#### MEG Data Processing

##### BESA Analyses

MEG data were first processed using BESA Research 6.1 (MEGIS Software GmbH, Grafelfing, Germany). A two-step process was employed for removal of muscle and movement artifact, with all analyses done blind to diagnosis. First, the participant’s raw EOG and MEG data were visually examined to remove epochs with blinks, saccades, or other significant EOG activity from the MEG data. Second, each participant’s MEG data were visually inspected for muscle-related activity, with a focus on data from sensors close to the temporalis muscles, and epochs containing muscle activity were removed. t-tests showed that groups had similar amounts of artifact-free RS data at Visit 1 (TD mean = 780 s, SD = 169; mTBI mean = 803, SD = 159; *p* > 0.05) and Visit 2 (TD mean = 774 s; SD = 206, mTBI mean = 806; SD = 155, *p* > 0.05).

For each RS recording, MEG and sMRI data were co-registered using BESA 6.1. In cases where sMRI data could not be collected due to scheduling conflicts (TD = 4, mTBI = 10), the participant’s MEG data were co-registered to an age-matched template (53). To constrain the scope of analyses (i.e., not examining group differences in many different frequency bands (e.g., delta, theta, alpha, beta, low gamma, high gamma)), an initial analysis determined the frequency where group differences were most pronounced at Visit 1. To this end, a standard BESA source model with 15 regional sources was applied to project each child’s raw MEG surface data into brain source space, where the waveforms are the modeled source activities (given two orthogonal dipoles per regional source, there are two time series at each location; Hoechstetter et al. (54)). These regional sources are not intended to correspond to precise neuroanatomical structures but rather to represent neural activity at coarsely defined regions and to provide measures of brain activity with better signal separation and with a greater signal-to-noise ratio than would be afforded at the sensor level (55, 56). The location of the regional sources in the model are such that there is an approximately equal distance between sources (3 cm), helping to separate signals originating from different brain regions.

To transform MEG data from the time domain to the frequency domain, a Fast Fourier Transform was applied to artifact-free 3.41 s epochs of continuous MEG sensor data to produce each of the two orthogonally oriented time series at each regional source. Each 3.41 s epoch overlapped 50% with the next epoch, and each epoch was multiplied by a Hanning window. This combination of overlap and windowing ensured that each time point contributed equally to the mean power spectra. When the segment of data following an artifact-free epoch was bad, there was no overlap between epochs, and windowing was thus applied to the next available artifact-free 3.41 second epoch. The mean power spectra for the two orthogonally oriented time series at each regional source were summed to yield the power at a given frequency at the source.

Group differences in RS power were examined from 4 to 56 Hz via uncorrected t-tests across all 15 regional sources, which identified beta-band frequencies (16–30 Hz) as the primary band showing group differences (see results). These 795 non-independent t-tests were done not as inferential tests about group differences but as a first characterization of the data in order to limit the number of tests in subsequent inferential analyses. For the subsequent whole-brain analyses, only a standard frequency band where groups showed the largest difference was further examined using Fast Vector-based Spatio-Temporal Analysis of L1-minumum (Fast-VESTAL, (57)).

##### Fast-VESTAL Analyses

For the Fast-VESTAL beta-band analyses, a participant-specific realistic boundary element method (BEM) head model was used for MEG forward calculation (58). The BEM mesh was constructed by tessellating the inner skull surface from the participant’s T1-weighted MRI into ∼6000 triangular elements with ∼5 mm edges. That grid was used for calculating the MEG gain (i.e., lead-field) matrix, which produced a grid with ∼7000 nodes covering cortical and subcortical gray matter.

For the Fast-VESTAL analyses, the eyes-closed time-domain data were filtered to examine beta (16–30 Hz) using a frequency-domain band-pass filter with zero phase shift via discrete Fourier transform. The MEG data were divided into 2.5 s epochs with 50% overlap. Fast-VESTAL was then used to obtain the source amplitude (root mean squared) MEG beta source images. In this approach, a fast Fourier transform was first performed for each epoch. This step transferred the time-domain MEG sensor-waveform signal into frequency-domain Fourier coefficients for different frequency bins within the beta band. Next, L1 minimum-norm-based frequency-domain Fast-VESTAL analysis was performed to obtain MEG source images for the real and imaginary part of the Fourier coefficients (59). The frequency-domain source power for each grid node was obtained by summing the power represented in the real and imaginary parts across all frequency bins within each band. The above procedure was performed for each epoch, and then the mean source power across all epochs was obtained for each grid node. The outcome of frequency-domain Fast-VESTAL analysis was a set of RS beta source power images for use in the group analysis (59).

For the Fast-VESTAL group analyses, the frequency-domain Fast-VESTAL source grid was obtained by sampling gray-matter areas from the participant’s T1-weighted MRI. Individual Fast-VESTAL source images for beta activity (16-30 Hz) were registered to Montreal Neurological Institute (MNI) space using an affine transformation (FLIRT–FMRIB’s Linear Image Registration Tool) in FSL (www.fmrib.ox.ac.uk/fsl/).

### Statistical Analyses

Independent-samples *t*-tests examined group age differences at Visit 1 and Visit 2. Fisher Exact tests assessed group differences in the proportion of males and females at each timepoint.

#### BESA Analyses

These initial analyses examined group differences in RS activity (4 to 56 Hz) at the 15 regional sources via t-tests, with the subsequent distributed source localization analyses assessing group differences in the frequency band in which groups showed the largest difference at Visit 1. As detailed in the Results, group differences were largest in the beta range.

#### Fast-VESTAL Analyses

To evaluate group differences in beta activity, a voxel-wise linear mixed-effect analysis was performed using the 3dLMEr package available in AFNI (https://afni.nimh.nih.gov/pub/dist/doc/htmldoc/programs/3dLMEr_sphx.html), with each adolescent’s whole-brain Fast-VESTAL beta image entered as the dependent variable, with group (mTBI and TD), Visit (Visit 1 and 2), and the Group X Visit interaction included as fixed effects and with subject entered as a random effect. Significant regions of interest (ROIs) that included ‘group’ were identified (i.e., an overall main effect of group, a simple main effect of group at Visit 1 or 2, or a Group x Visit interaction). Given that the focus of the study is differences between TD and mTBI adolescents, significant Visit effects not including Group are not reported. When a significant Group effect for an ROI was observed, an average beta value from that ROI was obtained by averaging the beta power across all voxels within the ROI to compute Cohen’s *d* effect sizes. For Fast-VESTAL group statistics, family-wise correction was determined using False Discovery Rate (FDR; (60)) as implemented in AFNI (3dFDR; https://afni.nimh.nih.gov/pub/dist/doc/htmldoc/statistics/fdr.html).

To examine associations between beta activity (the 3D Fast-VESTAL dataset) and symptoms (a 1D vector) in the mTBI group, voxel-wise correlations were performed using 3dTcorrelate in AFNI (https://afni.nimh.nih.gov/pub/dist/doc/program_help/3dTcorrelate.html). As shown in Table 1, in light of the considerable variability across adolescents with mTBI in the number of days between their concussion and their Visit 1, as well as variability in number of days between Visits 1 and 2, correlation were computed between Visit 2 minus Visit 1 difference in RS beta power and the Visit 2 minus Visit 1 difference in Teen PCSI Total score.

**Table 1.**
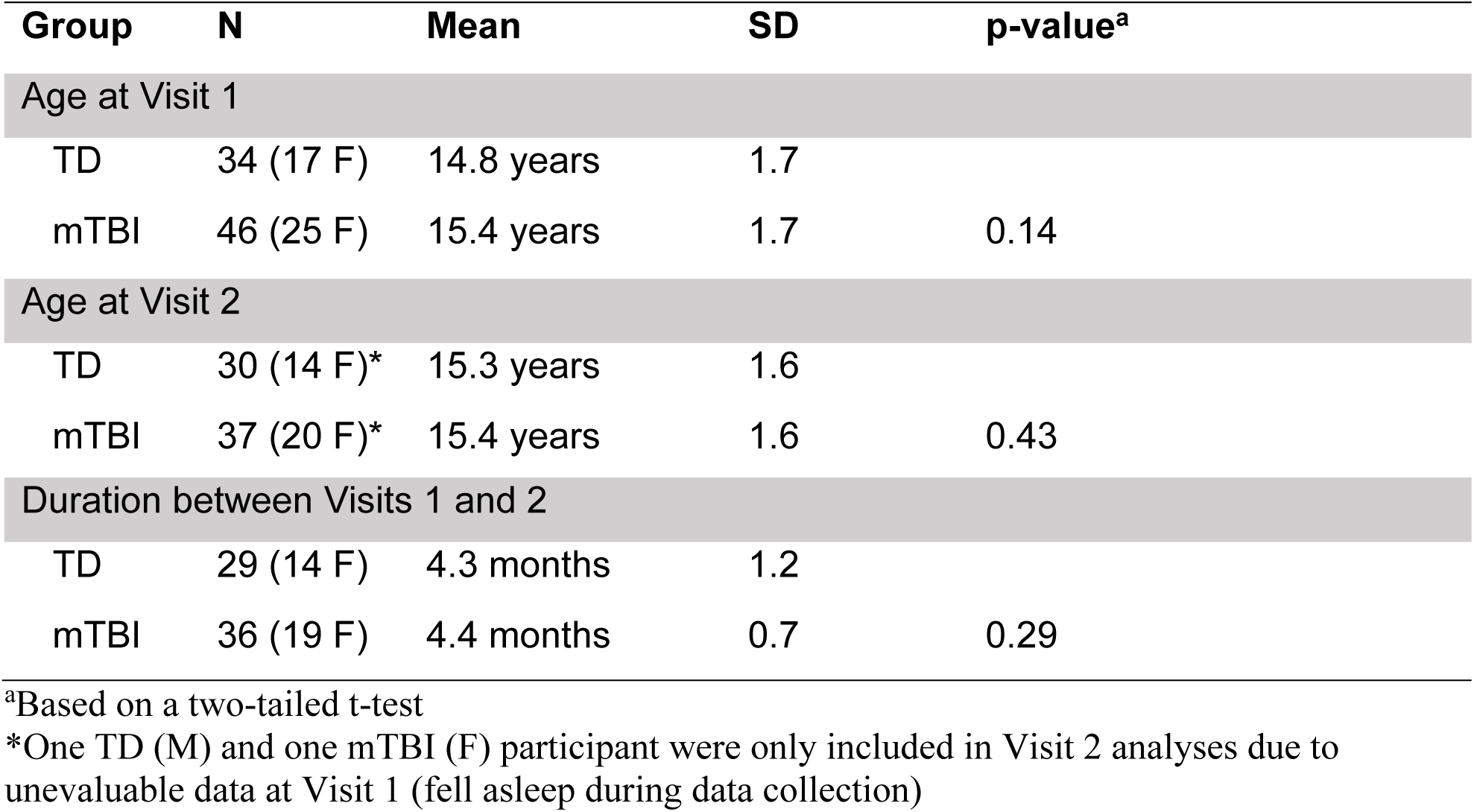

## Results

### Characteristic of the sample at each visit

Forty-six TD adolescents and 52 adolescents with mTBI passed the phone screen and were enrolled in the study. At Visit 1, 8 TD and 5 mTBI participants were excluded due to study psychologist concern for an anxiety and/or mood disorder based on the constellation of clinically significant elevations on self-report and parent-report questionnaires (e.g., YI-4R, CASI-5, CBCL, SCARED, CDI-2). Three TD adolescents were excluded from the study due to excessive metal artifact in the MEG recording (e.g., hair extensions or metal fillings). Finally, 2 participants (1 TD and 1 mTBI) were excluded from the study due to non-evaluable RS data at Visit 1 and 2, falling asleep during both exams. One TD and 1 mTBI participant were excluded at Visit 1 but included at Visit 2 due to falling asleep during the first but not the second visit recording. Given the above, evaluable Visit 1 MEG data were available from 34 TD adolescents and 46 adolescents with mTBI.

Reasons for a missed Visit 2 were as follows: COVID-19 pandemic lab closure (N = 3 mTBI), a new exclusion diagnosis at Visit 2 (N = 1 TD), family no longer wishing to participate, primarily due to COVID-19 concerns (N = 4 mTBI, 2 TD), family did not reply to emails and calls to schedule a second visit (N = 3 mTBI, 1 TD), or placement of braces after Visit 1 (1 TD). Given the above, evaluable Visit 2 MEG data were available from 29 TD adolescents and 36 adolescents with mTBI. As shown in Table 1, the groups did not differ in age at Visit 1 or 2 or time between visits. Fisher’s Exact tests showed no group differences in sex at Visit 1 (*p* = 0.82) or Visit 2 (*p* = 0.36).

Table 2 reports time between the mTBI and Visit 1 for the mTBI group and time between Visit 1 and Visit 2 by group and sex. Visit 1 data for the mTBI group were obtained on average 31 days after their mTBI injury (SD = 25 days, range 4 to 98 days). Visit 2 data were obtained on average 4._b_3 months (SD = 1.2 months) after their mTBI and on average 4.4 months (SD = 0.7 months) after Visit 1 for the TD group.

**Table 2.**

| **mTBI Time Between Concussion & Exam** | **# of Males** | **# of Females** |
| --- | --- | --- |
| Within 2 weeks | 8 | 8 |
| 2-4 weeks | 8 | 3 |
| 1 to 2 months | 2 | 10 |
| 2 to 3 months | 3 | 3 |
| 3 to 4 months | 0 | 1 |

| **mTBI Time Between First and Second Exam** | **# of Males** | **# of Females** |
| --- | --- | --- |
| Within 4 months | 6 | 7 |
| 4 to 5 months | 9 | 11 |
| 5 to 6 months | 1 | 0 |
| 6 to 7 months | 2 | 1 |

| **Controls Time Between First and Second Exam** | **# of Males** | **# of Females** |
| --- | --- | --- |
| Within 4 months | 7 | 9 |
| 4 to 5 months | 7 | 5 |
| 5 to 6 months | 0 | 0 |
| 6 to 7 months | 2 | 0 |

Online Supplement Table 1 provides details about the head trauma event for each mTBI participant. The injury was due to the following: 72% sports-related (e.g., football, lacrosse, basketball), 13% other such head impact (e.g., bumped into steel beam, fall), 11% motor vehicle crash, and 4% assault. A sports-related mTBI was more common in males (19 of 21) than females (15 of 26; Fisher’s exact test *p* < 0.05).

Paired-t-tests showed that PCSI total scores in the mTBI participants decreased from Visit 1 (mean = 29.93, SD = 21.29) to Visit 2 (mean = 15.33, SD = 17.05; *p* < 0.001), indicating clinical improvement. Of note is that 4 of the mTBI participants who were 12 years old were administered the Child version of the PCSI. Of these 4, 2 were administered the Child version at Visit 1 and 2, and 2 the Child version at Visit 1 and the Teen version at Visit 2 (they turned 13 after Visit 1). Given differences between the Child and Teen versions of the PCSI (e.g., very different range of scores), in these 4 adolescents it was not possible to assess changes in their mTBI symptoms or to include them in the analyses examining associations between brain function and concussion symptoms.

### MRI Findings

All MRI exams were read as normal by a pediatric neuroradiologist. Seven adolescents with mTBI and 6 TD adolescents were found to have increased FLAIR or T2 signal, the findings judged as nonspecific or unrelated to concussion.

### Visit 1 Group Differences in Regional Source Strength: 4 to 56 Hz

As shown in Figure 2, at Visit 1 more 4 to 56 Hz activity was observed in the TD group (blue mean line and 95% CI) than the mTBI group (red mean line and 95% CI) in several brain regions, with group differences most prominent in frontal and central regions. Group differences were most pronounced in the beta band (∼16 to 30 Hz) as indicated by the smallest *p*-values in the beta range (e.g., \**p* < 0.001 in light blue versus \**p* < 0.01 in magenta). Figure 3 shows the beta group differences (TD > mTBI) at the Midline Central location. Given that at Visit 1 group differences were largest in the beta range, the Fast-VESTAL whole-brain LMM analyses (including Visit 1 and 2) examined beta group differences.

**Figure 2.**
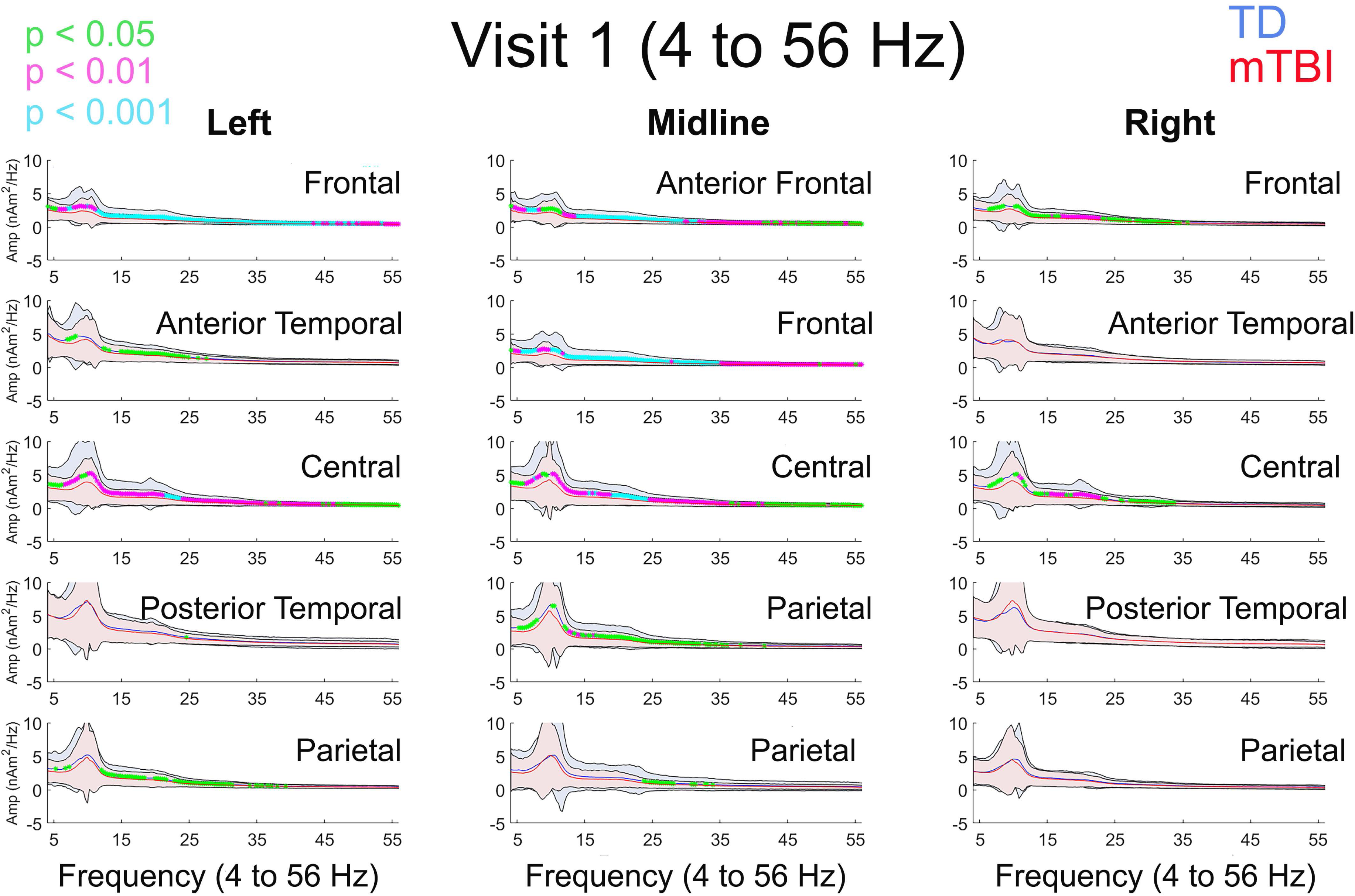
t-tests compared TD (blue mean line and 95% CI) and mTBI (red mean line and 95% CI) at Visit 1 at each of the BESA 15 source-model locations from 4 to 56 Hz (x axis). Across the examined frequencies, more RS activity was observed in TD than mTBI, with group differences most prominent in frontal and central regions, and with group differences most pronounced in the beta-band (∼16 to 30 Hz; highlighted in red circles) as indicated by the smallest *p*-values in the beta range (e.g., \**p* < 0.001 in light blue versus \**p* < 0.01 in magenta).

**Figure 3.**
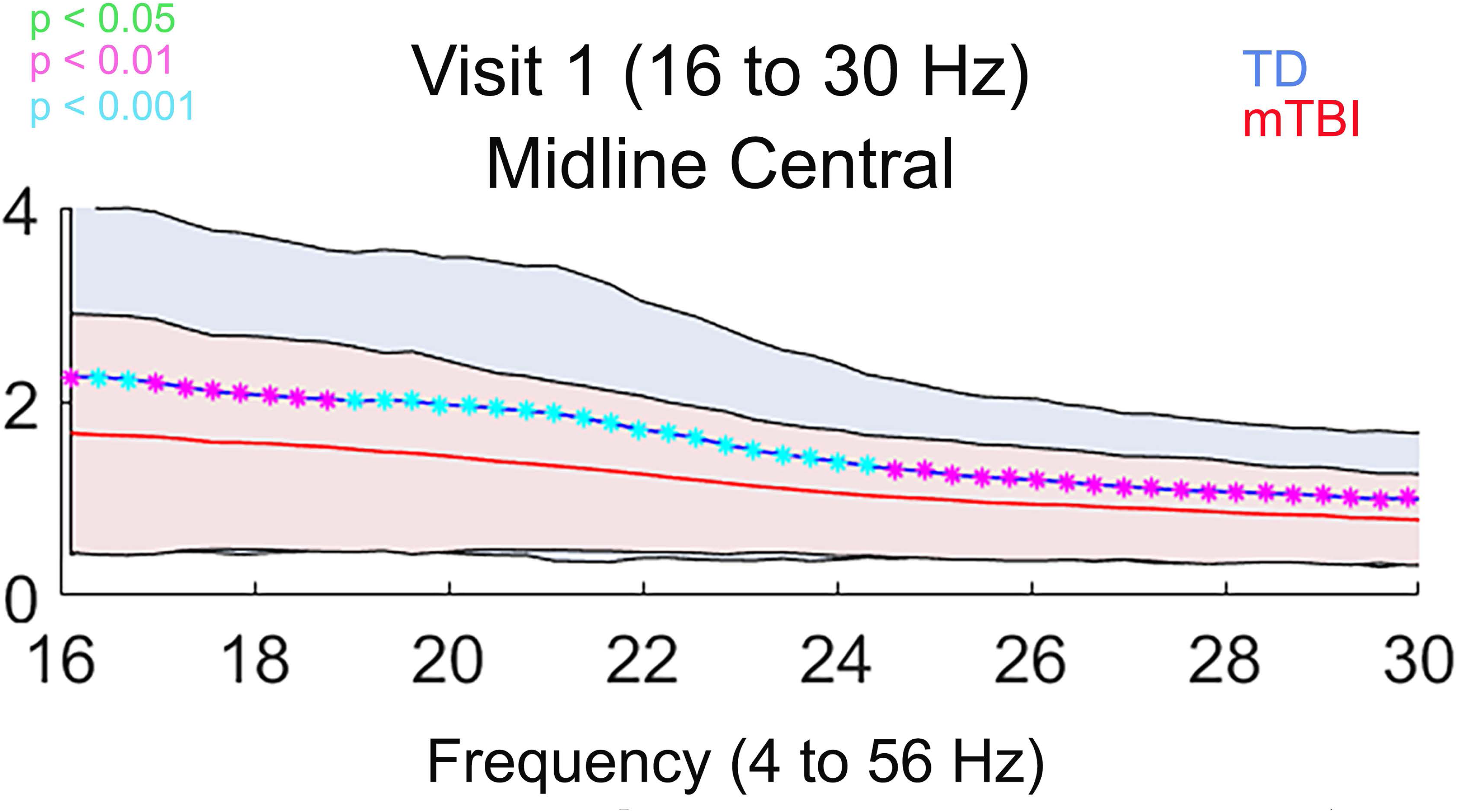
More beta activity in TD (blue mean line and 95% CI) than mTBI (red mean line and 95% CI) shown for the Midline Central location. Given that group differences were largest in the beta range (\**p* < 0.001 in light blue versus \**p* < 0.01 in magenta), the Fast-VESTAL whole-brain analyses examined beta (16 to 30 Hz) group differences.

### Whole-Brain analyses: 16 to 30 Hz

The Figure 4 left column shows the significant Group effects (across Visit 1 and 2) from the LMM analysis. To help interpret the findings, the Figure 4 right panel shows the group beta power means (with standard error bars) for Visit 1 and 2 for each ROI.

**Figure 4.**
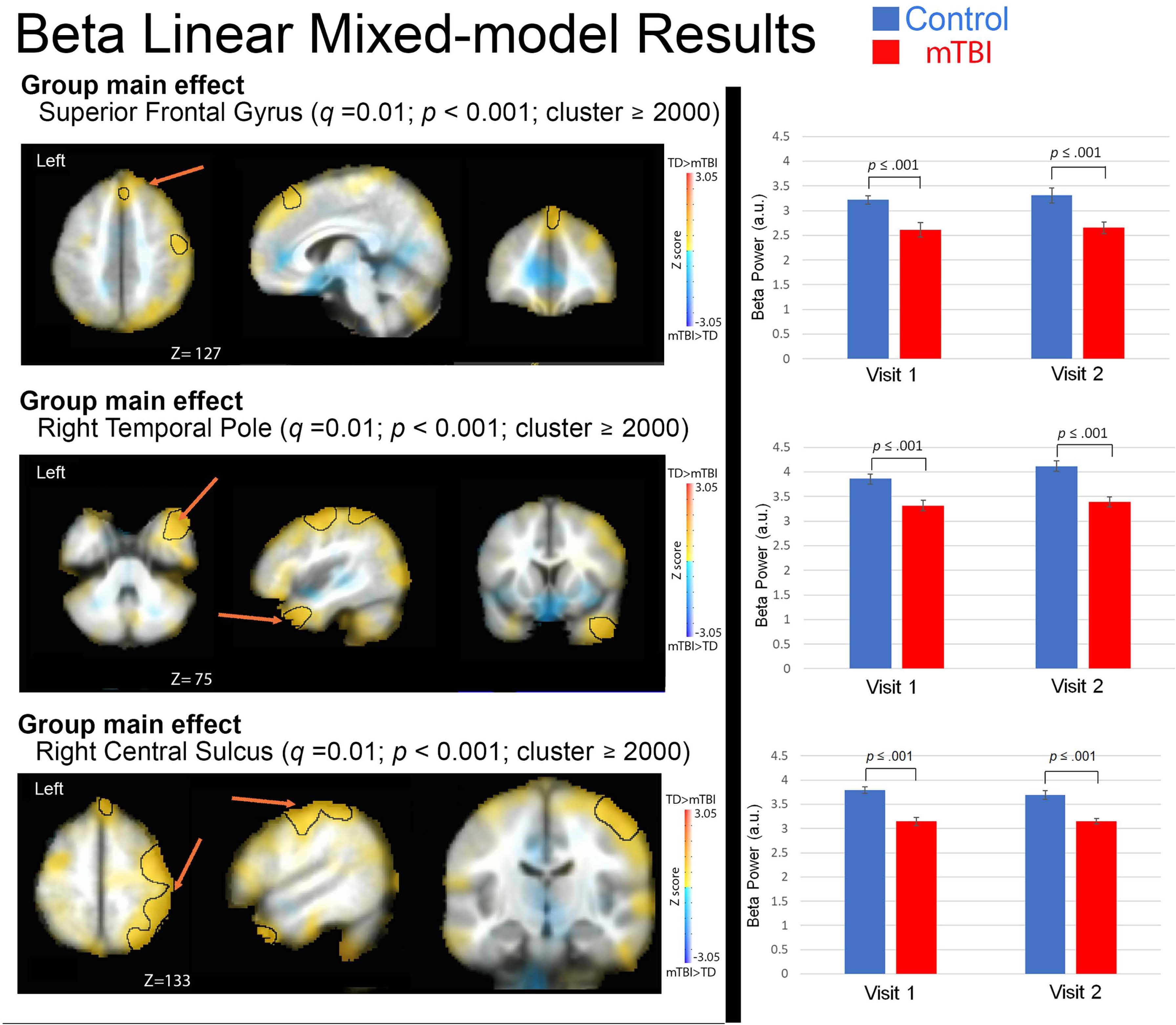
Left panel: group effects from the linear mixed-model analysis with family-wise correction (as implemented in AFNI). ROIs exceeding family-wise correction are circled in black (TD > mTBI in all 3 ROIs), and with brain areas showing non-significant TD>mTBI activity (shown in yellow) and mTBI>TD activity (shown in blue) displayed to better show the regional specificity of each group main effect. Right panel: for each ROI the TD (blue) and mTBI (red) group means (with standard error bars) at Visit 1 and Visit 2.

As shown in Figure 4, significant main effects of Group (TD > mTBI, *q* = 0.01) were observed at superior frontal gyrus, right temporal pole, and right central sulcus, No Visit main effects or Group x Visit effects were observed. Table 3 reports Cohen’s *d* effect sizes for the three Group ROIs, for the full sample as well as for males and females.

**Table 3.**

| Region of Interest | Cohen's $d^*$ | | | | | |
| --- | --- | --- | --- | --- | --- | --- |
|  | Full Sample |  | Males |  | Females |  |
|  | Visit 1<br>(n=80) | Visit 2<br>(n=67) | Visit 1<br>(n=38) | Visit 2<br>(n=33) | Visit 1<br>(n=42) | Visit 2<br>(n=34) |
| Group effect: Superior Frontal Gyrus | 0.75 | 0.84 | 0.70 | 0.98 | 0.78 | 0.72 |
| Group effect: Temporal Pole | 0.83 | 1.15 | 1.58 | 1.06 | 0.27 | 1.25 |
| Group effect: Right Central Sulcus | 1.30 | 1.31 | 1.48 | 1.08 | 1.19 | 1.80 |

\*A positive Cohen's $d$ indicates mTBI < TD

Finally, although sample size precluded including sex as a factor in the statistical model, Online Supplement Figure 1 shows for each ROI the group means and standard error bars as a function of Visit and Sex to allow readers to make qualitative comparisons and as a reference for future studies. In Supplement Figure 1, the left panel reproduces the left panel of Figure 4 and the right panel separates the right panel of Figure 4 by sex. Very similar patterns are evident for males and females.

### Beta-Power Associations with Symptoms

Associations between the change in concussion symptoms (Visit 2 minus Visit 1 PCSI total score) and change in beta power (Visit 2 minus Visit 1 Fast-VESTAL beta maps) were assessed. Figure 5 shows the findings for the Teen PCSI Total Score analyses. In the scatterplots, a negative PCSI change score (x axis; Visit 2 minus Visit 1) indicates fewer symptoms at Visit 2 than Visit 1 (i.e., clinical improvement), and a positive beta change score (y axis; Visit 2 minus Visit 1) indicates more beta-band activity at Visit 2 than Visit 1. Four ROIs were identified (*p* < 0.005, cluster = 1000), with Figure 5 showing a negative association (clinical improvement associated with increased beta power) in (1) left middle frontal gyrus, (2) left superior angular gyrus, and (3) right temporal parietal-occipital region and showing a positive association in (4) right temporal pole (clinical improvement associated with decreased beta power).

**Figure 5.**
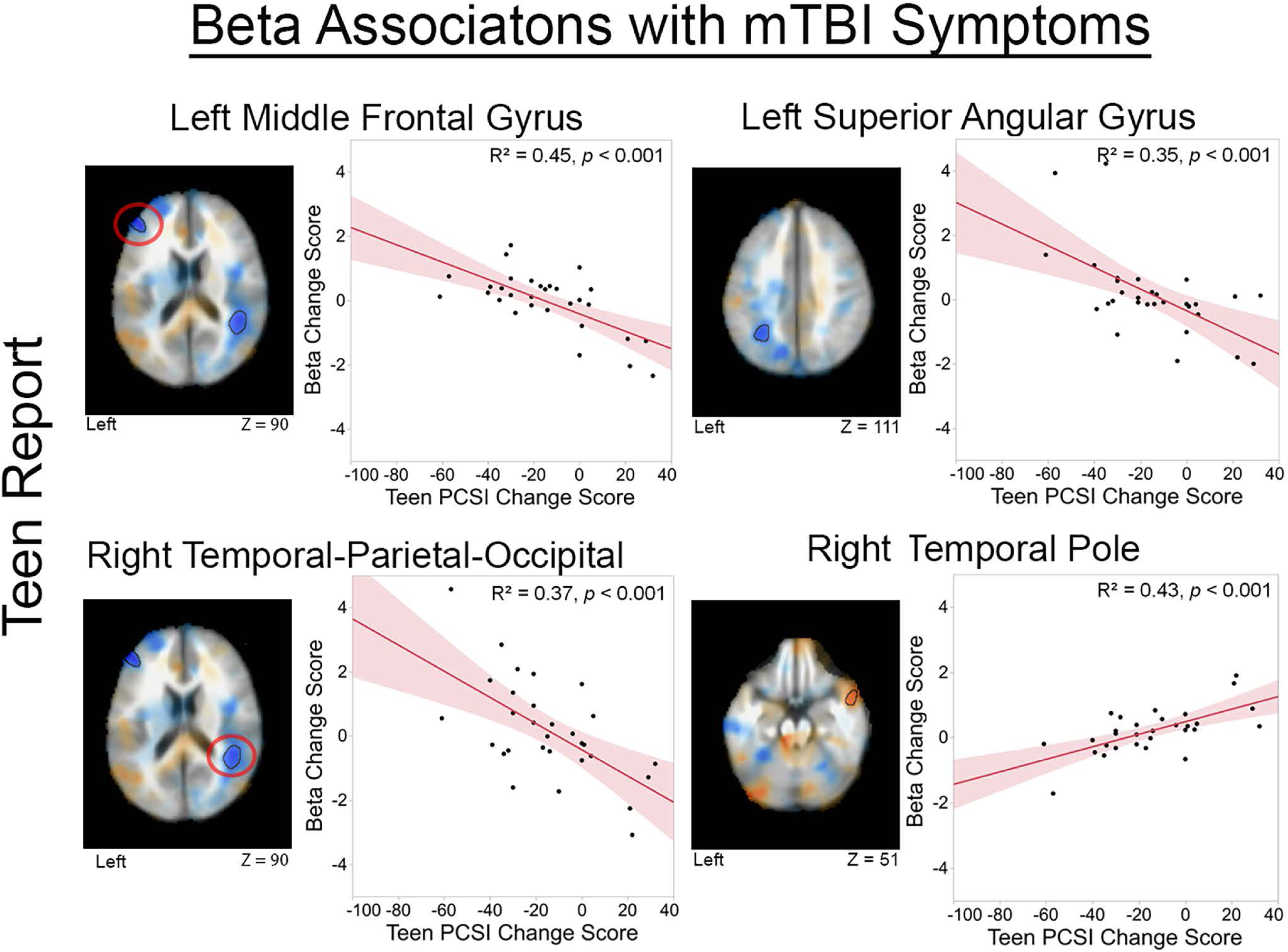
ROIs (family-wise corrected) showing associations between the Visit 1 to 2 change in mTBI symptoms (x axis; Teen PCSI total score) and Visit change in beta activity (y axis; Visit 2 minus 1 Fast-VESTAL beta maps). In the scatterplots, a negative PCSI change score (x axis; Visit 2 minus Visit 1) indicates fewer symptoms at Visit 2 than Visit 1 (i.e., clinical improvement), and a positive beta change score (y axis; Visit 2 minus Visit 1) indicates more beta-band activity at Visit 2 than Visit 1.

## Discussion

Consistent with previous studies (15–19), the clinical MRI exam did not identify structural brain abnormalities specific to the mTBI in a diagnostically clean (no previous DSM diagnosis) sample of male and female adolescents with mTBI. In contrast, MEG source analysis revealed regionally specific RS beta-band abnormalities. At Visit 1 (sub-acute period, 0 to 3.99 months post-concussion) and Visit 2 (sub-chronic period, 4+ months post-concussion), adolescents with mTBI showed less beta power at right central sulcus, right temporal pole, and bilateral medial superior frontal gyrus ROIs. Furthermore, RS beta power was observed to be of clinical significance, in that decrease in mTBI symptoms was associated with beta power increase from Visit 1 to Visit 2 in several brain regions. These findings suggest considerable potential clinical value of this assessment of brain function in the course of mTBI.

### Resting-State Abnormalities in Adolescents with mTBI

The finding of less RS beta activity in adolescents with mTBI (see Figure 4 and Table 3) extends that of Zhang et al. (31), who reported the same finding in adult males with mTBI (their MEG RS data obtained within 3 months of injury). In both studies, group differences were most prominent at areas surrounding central sulcus and frontal regions. In both studies, central sulcus group differences were more prominent in the right than left hemisphere. These two studies suggest RS beta activity as a measure for use in adolescents and adults within the first few months following a mTBI. Notably, these studies analyzed RS MEG using different distributed source localization methods, evidence that the ability to detect group differences is robust to source localization method.

With respect to group differences observed in the present study, two details are of note. First, whereas the right central sulcus group difference (across Visit 1 and 2) suggests atypical brain activity in primary sensory and motor areas, the other two group differences were observed in transmodal areas, brain regions that receive input from multiple sensory areas and that are associated with more complex or integrated cognitive activities (61). As reviewed in Zhang et al. (31), beta activity is associated with top-down attention (62), sensorimotor integration (63, 64), and motor planning and error monitoring (65). They noted that it is likely that axonal injury (66, 67), as well as disruptions to glutamate and GABA and thus a dysregulated excitation-inhibition balance (68), contributes to the abnormal beta activity in mTBI. The present and Zhang et al. findings suggest that primary motor and somatosensory areas are especially vulnerable to mTBI. Zhang et al. hypothesized that the cortical beta abnormalities observed in mTBI are due to damage to thalamocortical pathways. In particular, as lower frequencies (30 Hz and below, where most of the present findings occurred) are thought to reflect descending/top-down information processing generated in the deep infragranular layers, such findings would be driven by strong and reciprocal thalamocortical and subcortical connections (69). The finding of attenuated beta activity in mTBI is consistent with abnormal corticothalamic connectivity. Finally, the present finding of group differences in bilateral medial superior frontal gyrus is consistent with microscopic- and molecular-level damage to frontal areas being common after a mTBI (70, 71). Observing abnormalities in primary sensory/motor areas as well as frontal transmodal areas is intriguing, given that the principal gradient of functional connectivity tracks a functional hierarchy from primary sensory processing to higher-order functions, with damage in one area likely impacting neural activity in functionally connected areas (72, 73).

Not all present adolescent findings align with the MEG studies of mTBI in adults. As an example, in the present study, the BESA 15-source analyses identified theta, alpha, and gamma RS group differences (mTBI < TD; see Figure 2). These gamma findings are opposite those of Huang et al. (74), who observed more RS gamma in adults with blast-related mTBI and persistent post-concussive symptoms than controls. Huang et al. (74) hypothesized that that may reflect chronic dysregulation of gray-matter inhibitory interneurons and pyramidal cell networks. It may be that, if the mTBI adolescents in this study were followed longer post-injury, those with persistent post-concussive symptoms would eventually present with abnormally high beta and gamma activity. It may also be, however, that the Huang et al. findings are specific to a head injury due to a blast-related mTBI and thus not observed in the adolescents with injuries from other mechanisms.

To our knowledge, only 4 MEG studies have examined RS activity in adolescents with a mTBI, 3 with rather small samples. Building on the Lewine et al. (23, 24) mTBI adult delta-band findings, Davenport et al. (38) examined the pre- to -post-season change in RS delta-band activity (three minutes of artifact-free eyes-open data) in eight male high-school football players diagnosed with a concussion during the season, eight male high-school football players without a concussion, and eight male non-contact controls. They found that the football players with concussion had more delta-band activity at their postseason scan than the non-concussed football players and the noncontact controls. In the present study, the BESA 15 regional source analyses showed decreased delta activity in the mTBI cohort than in the TD participants. Key differences that make comparisons between the two studies difficult include the fact that Davenport et al. compared brain activity in the mTBI group before and after their injury. It is possible that in the present study similar delta-band findings would have been observed in the adolescents with mTBI comparing brain activity before and after the mTBI. Davenport et al. also examined only whole-brain relative delta power and did not assess region-specific effects or group differences.

Huang et al. (40) reported group differences in children and adolescents as a function of brain region. They examined RS activity in children and adolescents 8 to 15 years old (12 mTBI (83% males) and 12 controls (92% males)), with an average of 6 months between injury and RS exam. Huang et al. observed group differences across all examined frequency bands (delta-theta to gamma). As in the present study, the pattern of findings differed as a function of brain region. For example, the adolescents with mTBI showed beta hyperactivity in right temporal pole/anterior insular, left precuneus, and left cerebellum versus beta hypoactivity in bilateral dorsolateral prefrontal cortex, left ventromedial prefrontal cortex, and bilateral inferior temporal lobe. Differences between Huang et al. and the present study that might account for the differences in beta findings are many, including Huang et al. obtaining RS MEG measures on average 6 months post mTBI and examining mostly males. However, both studies indicate that the pattern of group differences differs across brain areas.

Safar et al. (37) reported on RS neural activity (5-minute eyes-open task; regional power and brain-wide functional connectivity examined) in 16 children and adolescents (mean age 12.5 years, 11 males) with chronic mTBI. They found that delta power was higher in bilateral occipital brain regions and right inferior temporal gyrus in mTBI participants than in controls. Group differences in functional connectivity were also observed (e.g., less delta-band functional connectivity in mTBI), with multivariate classification analysis showing higher group classification accuracy using functional connectivity than using regional power measures. A significant difference between Safar et al. and the present study is that Safar et al. studied children with a chronic mTBI, with the average time from injury to the MEG recording 455 days (range 79-1557), with the majority of the children reporting at least one post-concussive symptom and with 69% reporting greater than 2 post-concussive symptoms.

Finally, Huang et al. (39) examined RS MEG in children and adolescents 8 to 15 years old, recruiting a cohort with mTBI (N = 59, 66% male, ∼3 weeks post-injury) and a cohort with an orthopedic injury (N = 39, 64% male). Focusing on delta and gamma activity, a machine learning approach was used to classify the two groups and to predict symptoms at 3 months. They found that a combined delta-gamma model predicted group status (area under the curve value 98.5%) and showed that the 3-week MEG measures were good predictors of the 3-week to 3-month change in symptoms. Similar to the present study, the pattern of group differences was region-specific. For example, they observed more RS delta (e.g., frontal poles, rostral anterior cingulate cortex), more gamma (e.g., frontal poles, inferior frontal gyrus, superior central areas), and less gamma (e.g., medial frontal cortices, interior temporal gyrus, lateral occipital cortex) in mTBI than the orthopedic injury group. Huang et al. findings and the present findings demonstrate the need to assess neural activity in brain space rather than anatomically ambiguous sensor space to understand brain injury in mTBI.

### Associations between Beta Activity and mTBI Symptoms

The observed associations between MEG activity and mTBI symptoms are evidence of clinical relevance for MEG assessment. Specifically, present data suggest a mechanistic link between brain injury, brain electrophysiology, and behavioral symptoms. Given variability in the timing of the visits from injury and between Visit 1 and Visit 2, present analyses assessing associations between RS beta activity and clinical symptoms examined change scores. Given likely between-individual variability in how a participant reports the severity of mTBI symptoms, this approach used each participant as their own control. It was expected that, given lower beta power in mTBI participants than the TD cohort at Visit 1, an across-time normalization of (increase in) RS beta power would be associated with fewer mTBI symptoms. As shown in Figure 5, this association was observed in several brain areas^1^. Of note, none of the identified ROIs included primary sensory areas, suggesting that normalization of long-range functional connectivity is associated with improvement in mTBI symptoms. Also of note is that in the right temporal pole a decrease in the PCSI Total score between Visits was associated with a decrease in beta activity between Visits. These findings were unexpected and need replication. As detailed in Allen et al. (25), although some studies have reported associations between RS activity such as increased low-frequency activity and mTBI symptoms scores (24, 59), other studies have not (23, 75). It is hypothesized that, given the regional specificity of mTBI findings, the pattern of findings is likely a function of the participants’ age and time since injury and also potentially of source-localization methods. In the present study, the longitudinal design provided the ability to use each participant as their own control, thus providing some control for the between-subject differences in the time of their Visit 1 and 2.

## Limitations

There are several limitations. First, recruitment (Visit 1) and follow-up (Visit 2) were significantly disrupted due to the COVID-19 pandemic, with the lab closed for several months early in the pandemic, and with many families delaying or canceling their visit after the lab reopened due to concerns about COVID-19. As a result, the exams did not always fall within the planned timeframe. Given this variability, associations between brain function and clinical symptoms were assessed via RS beta change scores and PCSI change scores, so that for each adolescent the RS and the PCSI change scores spanned the same time interval.

Second, although group differences were observed from 4 to 56 Hz, the present whole-brain analysis focused only on the frequency band where group differences were the most prominent, in order to limit the number of analyses. As noted throughout this manuscript, mTBI and TD group differences have been reported for lower and higher frequencies (for review see Allen et al. (25)). Other studies have used machine learning to combine RS measures to obtain an optimal differentiation of groups (e.g., (76)). Further investigation is necessary, including analyses examining RS functional connectivity (e.g., (29, 33)) and the temporal dynamics of the RS beta rhythms (e.g., (77)).

Fourth, given the sample size, biological sex was not included as a factor in the statistical analyses, although the sample was better balanced than most. A qualitative examination of the results indicated a similar pattern of findings for males and females. Assessing sex differences is of interest given brain injury sex effects in rodent models (e.g., female rats more likely than male rats to exhibit behavior deficits following a mTBI (78)) and similarly in humans (79–81).

Finally, to compare present findings to all previous MEG RS mTBI studies, RS activity was examined in a ‘standard’ frequency band, in this case the beta band. Several recent papers note limitations with this approach, in particular the need to consider RS activity as composed of periodic (RS oscillatory activity) + aperiodic activity (background brain activity) (82–84). Present analyses did not undertake that distinction, maximizing comparability with existing child and adult MEG RS studies, none of which did so.

## Conclusions

Evidence of regionally specific RS beta-band abnormalities (less beta-band power in mTBI than TD cohorts) was observed several months post-injury in a diagnostically clean sample of male and female adolescents with mTBI. The group effects were large, with Cohen’s d’s ranging from 0.75 to 1.31. Furthermore, RS beta increase in several brain regions from Visit 1 to Visit 2 was associated with decrease in mTBI symptoms Visit 1 to Visit 2. Present findings suggest that RS beta power has potential as a measure and perhaps as a mechanism of clinical recovery in adolescents with mTBI.

## Ethics Approval and Consent to Participate

This study was approved by the local Institutional Review Board, and all families gave written informed consent. All adolescents 12 to 17 years gave written assent to participate.

## Availability of Data and Materials

The datasets generated and/or analyzed during the current study will be made available upon completion of this study and reasonable request.

## Footnote

1. Analyses examining the brain function and clinical symptom association at the ROIs where group differences were observed are difficult to interpret given an expected regression to the mean in brain activity as well as mTBI symptoms across time. In particular, given that the group-difference ROIs identify the brain areas showing the largest group differences at Visit 1, across time the brain activity in individuals with mTBI is likely to move more towards the mean TD value. Concussion symptoms will on average also decrease as a function of time, this resulting in a brain activity and symptom correlation at the group-difference ROIs that is difficult to interpret.

## Acknowledgements

DISCLAIMER: Dr. Berman reports a consultancy with McGowan Associates. Dr. Roberts declares his position on the advisory boards of (a) CTF MEG, (b) Ricoh, (c) Spago Nano Medical, and (d) Prism Clinical Imaging. Dr. Roberts and Dr. Berman declare intellectual property relating to the potential use of MRI measures for detecting TBI. This study was supported by the Pennsylvania Department of Health Cure Award (4100077078) as well as NIH grant R01MH107506 (JCE) and NICHD grant R01HD093776 (JCE).

