## Supplementary figures and images for "Regionally specific resting-state beta neural power predicts brain injury and symptom recovery in adolescents with concussion: a longitudinal study"

### Supplemental Figure 1

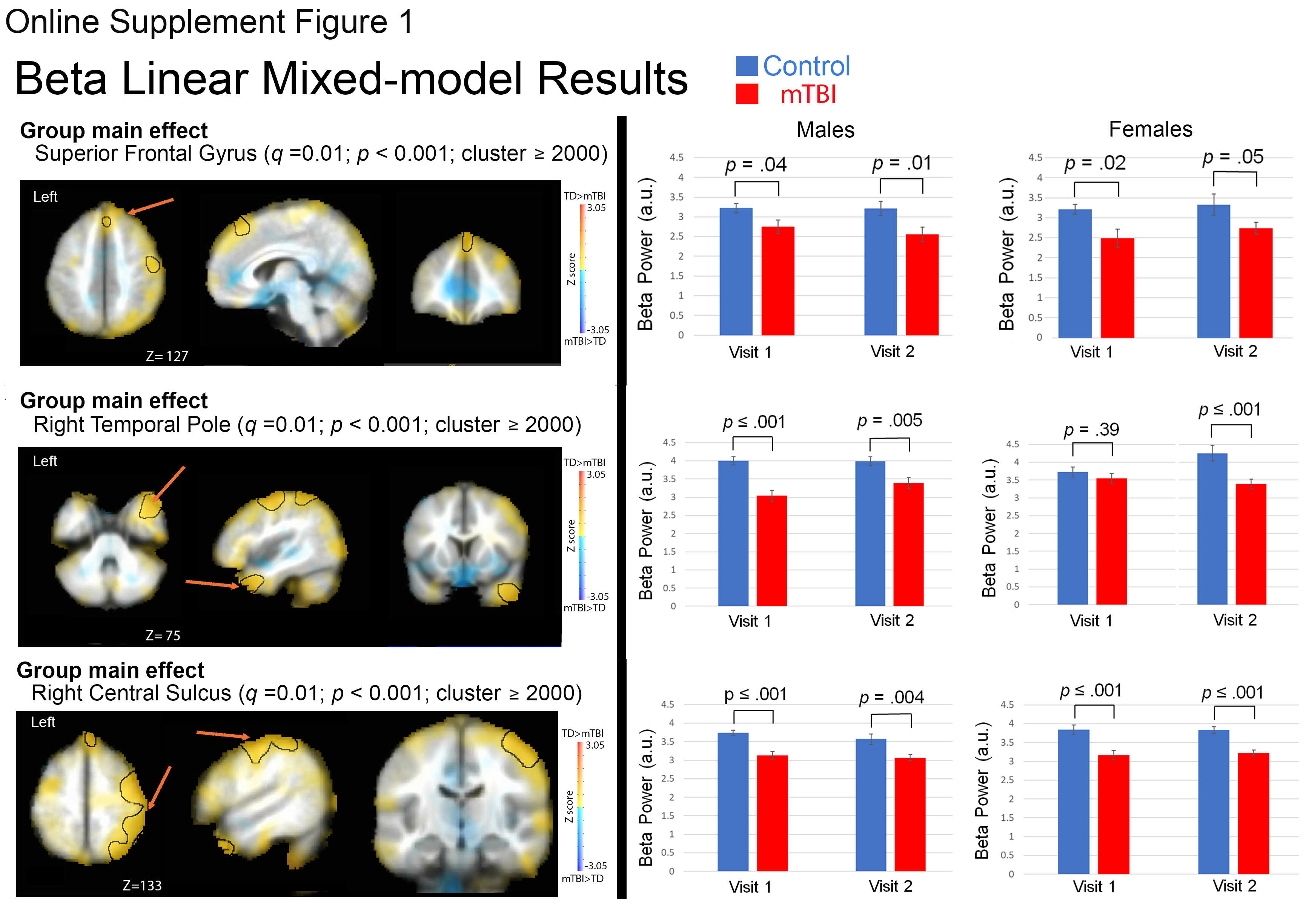

### Supplemental Table 1

Online Supplement Table 1

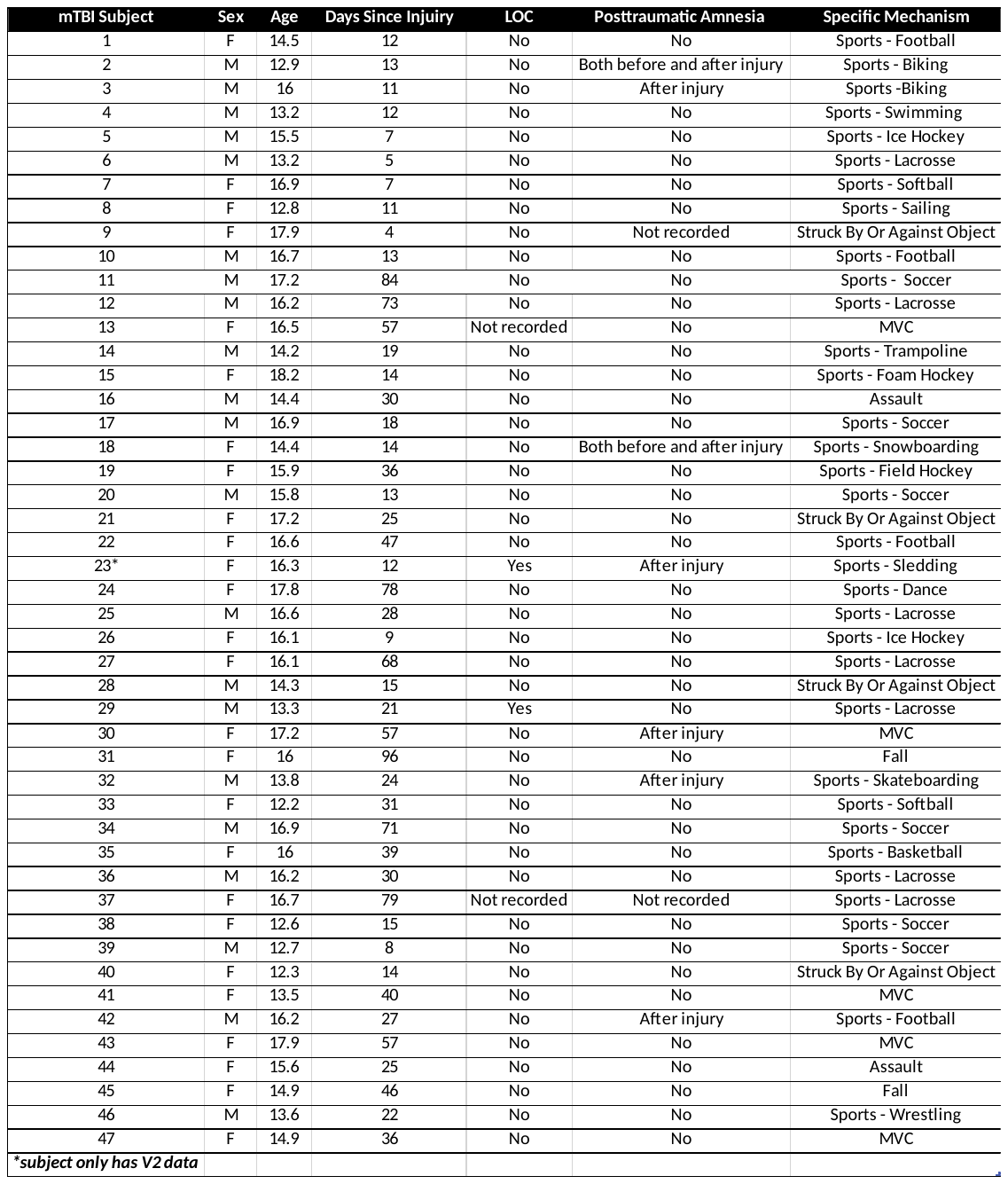
